# The dual activity of BBK32: Implications for simultaneous inhibition of borrelial-specific antibody-dependent complement activation and fibronectin binding

**DOI:** 10.1101/2024.06.07.598020

**Authors:** Alexandra D. Powell-Pierce, Charles E. Booth, Payton Smith, Brittany L. Shapiro, Shannon S. Allen, Brandon L. Garcia, Jon T. Skare

## Abstract

Complement inhibition is exploited by extracellular pathogens to combat clearance. *Borreliella burgdorferi*, the causative agent of Lyme disease, harnesses complement evasion techniques to establish and maintain infection in mammalian hosts. *B. burgdorferi* encodes *bbk32*, a surface lipoprotein that binds extracellular matrix (ECM) components, specifically glycosoaminoglycans (GAGs) and fibronectin (Fn) within its amino terminus. In addition to its ECM-binding functions, the carboxy terminal half of BBK32 binds to the C1r protease and prevents complement activation. This classical complement inhibitory activity protects *B. burgdorferi* from complement-mediated killing following the addition of normal human serum. Herein we demonstrate that full length BBK32 binds both Fn and C1 concurrently, indicating that binding of these macromolecules do not sterically hinder their simultaneous interaction. Given the link of antibody dependence to the classical pathway, we tested how the presence of BBK32 would protect *B. burgdorferi* from antibody mediated, complement dependent killing. The presence of BBK32 provided protection against borrelial-specific antibody binding and concomitant complement activation *in vitro*. We also demonstrated, using both fluorescence microscopy and flow cytometry, that the presence of BBK32 was required for reduced C4 deposition on the surface of borrelial cells. This work demonstrates the potential for BBK32 to simultaneously bind to both C1r and Fn and contributes to the broader understanding of the ability of *B. burgdorferi* to evade antibody-dependent complement-mediated killing. We contend that these observations ostensibly provide *B. burgdorferi* with coincident dissemination and immune evasion activities needed for optimal survival during infection.

**AUTHOR SUMMARY:** Lyme disease, caused by *Borreliella burgdorferi* and other related species, is the most common arthropod-borne infection in the United States. As an extracellular pathogen, *B. burgdorferi* is exposed to the complement system--a soluble proteolytic cascade that clears invaders. Complement is defined by three pathways known as the alternative, lectin, and classical. The classical complement cascade is activated by the binding of antibodies to a foreign or damaged cell. For *B. burgdorferi*, the BBK32 surface protein is known to mute the classical pathway by binding and inhibiting the initiating protease C1r. However, no studies have addressed how BBK32 protects infectious *B. burgdorferi* from borrelial-specific antibody binding and clearance. Here we show that native BBK32 protects infectious *B. burgdorferi* from antibody-dependent, complement mediated killing. Given BBK32s other activity—namely its known adherence to fibronectin—we were interested to test if BBK32 could bind the C1 complex, which contains C1r, together with fibronectin. Our results suggest that these complex macromolecules can bind BBK32 simultaneously. These observations suggest that the dual activity of BBK32, namely fibronectin binding and C1r inhibition, are not mutually exclusive and contribute to *B. burgdorferi*’s ability to establish infection and evade antibody-based host clearance, respectively.

## INTRODUCTION

*Borreliella burgdorferi*, the causative agent of Lyme disease, is responsible for the largest incidence of vector-transmitted illness in the United States [1–4]. Transmitted by hard ticks of the *Ixodes* genus and predominantly dependent on small rodents as a reservoir, *B. burgdorferi* can adapt to these disparate environments through a diverse set of genes regulated by environmental cues that have not yet been fully determined [5,6]. Differentially regulated genes contribute to its ability to establish infection in the vastly diverse sites it colonizes between ticks and vertebrates [6–11], as well as escape the immune systems of organisms at distinct points in its life cycle [12,13]. During vertebrate infection, *B. burgdorferi* evades an onslaught of immune components, both natural and targeted, that seek to kill the spirochete [14–16]. Given that *B. burgdorferi* is largely considered an extracellular pathogen, one branch of immunity that contributes to the killing of *B. burgdorferi* during vertebrate infection is the complement cascade [17,18]. The complement system is a series of proteolytic reactions that function to recruit immune cells to the site of activation, tag the target cell with opsonins for phagocyte recognition, and create a lytic pore in the membrane of the target cell, termed the membrane attack complex (MAC) [19–22]. The classical pathway of complement is initiated by bound antibody and the resulting deposition of the C1 complex comprised of C1q, C1r, and C1s [20,21,23]. Binding of two or more IgG class immunoglobulins, or one IgM molecule to surface antigens, is sufficient to initiate a conformational change in the C1q subunit that leads to activation. Specifically, the C1r initiating protease autocleaves and then cleaves C1s. Activated C1s then modifies C2 and C4 downstream via partial proteolysis that results in release of C4a, subsequent opsonization of the target cell(s), and formation of the MAC [20,21]. During tick transmission of *B. burgdorferi* in mice, the presence of murine natural IgM—that are not directed toward any particular invader, including *B. burgdorferi*—minimizes the number of spirochetes in the tick midgut, presumably due to complement-mediated killing [24]. Nonetheless, serum from mice previously infected with *B. burgdorferi* is more effective at spirochete destruction than naïve serum, implying a role for both natural and directed antibody responses in reducing *B. burgdorferi* numbers at different stages during infection [24,25]. The ability of *B. burgdorferi* to quell the classical complement cascade, following antibody recognition, would seemingly result in extended survival and persistence in the face of both a non-specific and specific humoral response.

*B. burgdorferi* encodes for a wide range of complement inhibitors that function to prevent killing mediated by the complement cascade, with most of the prior work focused on the alternative pathway and, to a lesser extent, the classical pathway [26–28]. Resistance to the alternative pathways is mediated by numerous surface-exposed proteins designated as CRASP, OspE, Erp, or Csp proteins [29–35]. These proteins facilitate the binding of factor H to the surface of the borrelial cells thereby preventing the activation of C3 convertase and are produced within *B. burgdorferi* at different stages of the zoonotic cycle [36,37]. For the classical pathway, several borrelial proteins have been identified that inhibit activation. For example, p43, a protein of unknown identity that functions as a recruiter of C4 binding protein, inhibits both the classical and lectin pathway [38]. Separately, OspC recognizes C4b, inhibiting C3 convertase formation [39]. Although p43 or OspC do not prevent C4 cleavage and release of C4a, they do impair deposition of C3b, a crucial component of complement, preventing further downstream proteins in the classical cascade from becoming activated. BBK32, which is upregulated during vertebrate infection [36,37,40] and specifically inhibits the classical pathway by binding to C1r, can prevent activation of the classical pathway of complement [41–44].

Prior to its discovery as a complement inhibitory protein, BBK32 was characterized as a fibronectin (Fn) binding protein, and then for its ability to bind glycosaminoglycans (GAGs) [45–49]. These ECM interactions are thought to play a role in *B. burgdorferi*’s ability to extravasate through the vascular endothelium and disseminate throughout the infected host, mediating its ability to infect and colonize distal tissues [45,49,50]. During murine infection studies, a *bbk32* mutant exhibits attenuated infectivity at lower infection doses [45,50]; however, this phenotype has not yet been attributed to either BBK32’s ECM binding or its complement inhibitory activities. Additionally, the role of targeted antibodies in *B. burgdorferi* classical pathway mediated killing, broadly and in the scope of BBK32’s activity, has not yet been established. Because the classical pathway is dependent on antibody binding to the *B. burgdorferi* surface for its activity, BBK32-mediated inhibition may be important in both the initial stages of pathogen clearance when “natural” IgM recognize *B. burgdorferi* and later once a targeted antibody response has been developed [24,25]. Antibody binding can lead to a wide range of immune-mediated pathogen clearance, from phagocyte binding and phagocytosis to activation of lymphocytes [20,21]. Previously, we have shown that key residues of BBK32 interface with C1r’s substrate specificity site and occlude the active site of its serine protease domain [42]. When specific sites are mutated from their native amino acids to alanine, BBK32’s binding to C1r and classical complement inhibitory activity is lost [42].

Earlier studies focused on identifying key regions of BBK32 interfacing with Fn and GAG, which are likely important for adhesion to vasculature during the dissemination phase of Lyme disease [51,52]. Both the GAG and Fn binding domains map to the structurally disordered amino terminal half of BBK32 [44,46–49,53]. The known interactions between BBK32 and its ligands Fn, GAG, and C1r have been studied independently *in vitro* [44,46–49,53]. A prior study showed that BBK32’s Fn- and GAG-binding activity served to slow down the spirochetes in the vasculature, thus promoting extravasation [51,52]. Large domain deletions in BBK32, when analyzed in mouse vasculature using intravital microscopy, showed that BBK32 functions to first tether *B. burgdorferi* to the vasculature via its Fn interactions, functioning as a brake, while GAG-binding positions the cells optimally for extravasation through longer-lived interactions [52]. Despite this, the spatiotemporal role of each function during infection, in particular BBK32’s C1r-binding activity, remains unclear. Additionally, the ability of BBK32 to interact with ECM components and C1r or the entire C1 complex, either independently or simultaneously, has not been determined.

Here, we use structural modeling and biochemical analyses to show that BBK32 can simultaneously interact with Fn and C1. Using an *in vitro* model to demonstrate the antibody-dependence of complement killing in an infectious isolate of *B. burgdorferi*, we also demonstrate the ability of BBK32 to partially protect borrelial cells from this lethal event. As a result of these studies, we have determined that BBK32’s adhesin and C1r inhibitory activity may play a role not only during initial stages of infection, but its ability to impair classical complement-mediated clearance after a robust borrelial antibody response has been mounted. In addition, these studies have highlighted BBK32’s ability to interface with both Fn and C1 concurrently, indicating that the role of these functions during dissemination do not require prioritization by the spirochete.

## MATERIALS AND METHODS

### Bacterial Strains and Plasmids

*Borreliella burgdorferi* B31 strains ML23 and B31-A3-GFP [54,55] were grown to mid-log phase in BSK-II and 6% normal rabbit serum (Pel-Freez Biologicals, Rogers, AR). The *bbk32* mutant derivatives of ML23 and B31-A3, JS315 and GP100, respectively, are referred to in **Table 1**. B31-A3 and GP100 were cultured with gentamicin at 50 µg/ml under conditions that partially mimic the mammalian environment (5% CO_2_, 37°C, pH 6.8). Additional antibiotic resistance for JS315 and GP100 are indicated in Table 1. *B. burgdorferi* strains containing either pBBE22*luc*, pCD100, or pAP7 were grown with kanamycin at 300 µg/ml.

**Table 1.** *Borreliella burgdorferi* strains used for present study.

| **Table 1. *Borrelia burgdorferi* strains used for present study.** |  |  |
| --- | --- | --- |
| ***Borrelia burgdorferi* Strain** | **Description** | **Reference** |
| ML23-pBBE22/*luc* | Serum-sensitive, non-infectious *B. burgdorferi* B31 derivative strain lacking linear plasmid 25 with shuttle vector pBBE22 encoding *bbe22* and *B. burgdorferi* codon-optimized *luc* gene under the control of a strong borrelial promoter (*P<sub>flaB</sub>-luc*); kan<sup>R</sup>. | [50,56] |
| JS315- pBBE22/*luc* | *bbk32* mutant in ML23-pBBE22/*luc* background; strep<sup>R</sup>, kan<sup>R</sup>. | [50] |
| JS315-pCD100 | JS315-*luc* with *bbk32* under control of its native promoter in pBBE22/*luc*; strep<sup>R</sup> kan<sup>R</sup>. | This study |
| JS315-pAP7 | JS315-*luc* with *bbk32* R248A/K327A under control of its native promoter in pBBE22/*luc*; kan<sup>R</sup>. | This study |
| B31-A3-GFP | B31-A3 transformed with *gfp* under control of the borrelial | [55] |
|  | *flaB* promoter on the cp26 plasmid; gent<sup>R</sup>. |  |
| GP100 | *bbk32* mutant in B31A3-GFP background; gent<sup>R</sup>, strep<sup>R</sup>. | This study |
| GP100-pCD100 | GP100 with *bbk32* under control of its native promoter in pBBE22/*luc*; strep<sup>R</sup> kan<sup>R</sup>. | This study |
| GP100-pAP7 | GP100 with *bbk32* R248A/K327A under control of its native promoter in pBBE22/*luc*; kan<sup>R</sup>. | This study |

### Transformation of *B. burgdorferi*

Transformation of strains *B. burgdorferi* ML23 with the plasmid construct pAP7, and B31-A3-GFP with pCD100 and pAP7 was performed as previously described [42,45,57–59] The presence of plasmids pCD100 and pAP7 was selected in complete BSK-II media using kanamycin at a final concentration of 300 μg/ml. As is the norm, all borrelial transformants were screened for their compendium of plasmid DNA [54]. Only transformants that maintained the collection of plasmids found in their parental derivative were used.

### Far Western Overlay Analysis

Far Western overlays were carried out essentially as described [44,57], using *B. burgdorferi* strains ML23-pBBE22*luc* and B31-A3-GFP along with their respective *bbk32* mutant derivatives (JS315 and GP100, respectively; see **Table 1**). The *bbk32* mutant derivatives (JS315 and GP100) were transformed with either pCD100 or pAP7, which encode for native *bbk32* or the *bbk32*-R248A/K327A allele, respectively **(Table 1)**. Whole-cell lysates were generated for the borrelial strains, and 2.5x10^7^ whole cell equivalents were resolved by SDS-PAGE, and then transferred to PVDF membranes. Membranes were blocked overnight in 5% non-fat milk, washed, and then incubated with either (1) 20 µg human fibronectin (EMD Millipore); (2) 20 µg active human C1 complex (Complement Technologies); or (3) a combination of Fn and C1 complex overnight. Membranes were then washed and probed for either fibronectin or C1r using an anti-human Fn HRP conjugate (Santa Cruz Biotechnology), or an anti-human C1r HRP conjugate (Santa Cruz Biotechnology), respectively (both diluted 1:5,000). To confirm BBK32 production, a monoclonal antibody to the C-terminus of BBK32 was used at a 1:10,000 dilution, and the samples normalized to FlaB levels using an anti-borrelial FlaB monoclonal antibody (US Biological). Both monoclonal antibodies were detected using a goat anti-mouse immunoglobulin HRP conjugate diluted 1:10,000 (ThermoFisher Scientific). Blots were then visualized using the Western Lightning Plus-ECL (PerkinElmer).

### C1r and Fn Binding Studies

Surface plasmon resonance (SPR) assays were performed using immobilized BBK32 on a CMD200 sensor chip (Xantex Bioanalytics) via standard amine coupling using conditions previously described [42–44,57]. 20 nM Fn or 50 nM C1, as well as 20 nM Fn and 50 nM C1 together, were individually injected over the BBK32 biosensor. Experiments were performed with each cycle having an association time of 2 min, a dissociation time of 3 min, and two 1 min injections of regeneration buffer (0.1 M Glycine (pH 2.0), 2.5 M NaCl).

The ability of BBK32 to concurrently interact with Fn and C1 was determined based on the binding response just prior to the injection stop and calculated by subtracting the Fn curve from the co-injection curve. This was then compared to the curve of the injection of C1 alone.

### Antibody-Dependent Complement-Mediated Killing Assays

Antibody-dependent complement killing was assessed *in vitro* by diluting cells to a concentration of 5x10^6^ *B. burgdorferi* cells/ml in PBS, 0.5% BSA. Cells were then exposed to 175 ng rabbit anti-*B. burgdorferi* polyclonal antibody (Abcam) in addition to 20% NHS (Complement Technologies) for 30 minutes similar to recently published work [60]. Control groups were exposed to 175 ng rabbit isotype antibody (Abcam) coupled with 20% NHS to depict the antibody-dependent binding associated with complement activation, or 175 ng anti-*B. burgdorferi* antibody coupled with heat inactivated NHS to test whether the killing was purely antibody-dependent and assess the complement-dependence on borrelial death, respectively. Cells were then assessed for viability using a dark field microscope based on motility and membrane disruptions, as done in earlier published work [42,57]

### Fluorescent Microscopy and Colocalization Analysis

Colocalization of anti-*B. burgdorferi* antibodies and complement component C4c was determined using the *B. burgdorferi* B31-A3-GFP strain derivatives after washing the cells and incubating with 1 µg of a rabbit anti-*B. burgdorferi* polyclonal antibody (Abcam). C5 depleted human serum (Complement Technologies) was added to 20% as a source of complement. Subsequently, cells were incubated for 30 minutes with 0.5 µg of a mouse monoclonal antibody to C4c (Quidel). Cells were washed with PBS with 0.5% BSA, and then incubated for 30 minutes with 1 µg anti-rabbit IgG-Alexa Fluor 594 Plus to detect anti-*B. burgdorferi* rabbit antibodies, as well as 1 µg anti-mouse IgG-Cy5 to detect bound C4c antibodies. Cells were then fixed using 4% methanol-free paraformaldehyde and mounted onto a slide using Prolong Diamond Antifade Mountant (ThermoFisher). Cells were imaged using the Olympus Fluoview FV3000 confocal microscope, with equivalent exposure times for each fluorophore channel across samples.

### Flow Cytometry

The preparation for the analysis of the GFP expressing *B. burgdorferi* isolates by flow cytometry was identical to that described above for fluorescent microscopy with minor experimental modifications. Initially, *B. burgdorferi* cells were incubated with 1 µg of a rabbit anti-*B. burgdorferi* polyclonal antibody (Abcam). C5 depleted human serum (Complement Technologies), to a final concentration of 15% was added as a source of complement. Following fixation, the samples were incubated for 30 minutes with 1 µg of a mouse monoclonal antibody to C4c (Quidel). Cells were washed with PBS with 0.5% BSA and then incubated for 30 minutes with 1 mg anti-rabbit IgG-Alexa Fluor 594 Plus to detect total borrelial cells and 1 mg anti-mouse IgG-Cy5 to determine the B. burgdorferi with C4 deposited on their surface. Cells were washed with PBS and analyzed using the BD Fortessa X-20 within 24 hours of sample preparation. 10,000 events were read for each sample on the flow cytometer. Cells were characterized based on their individual fluorescent properties and analyzed with BD FACSDiva^TM^ Software. *B. burgdorferi* were also prepared following the same protocol described but without staining. In addition, borrelial cells were individually stained with each secondary antibody to establish proper gating and demonstrate primary antibody specificity.

### AlphaFold 2 analysis

AlphaFold2 (AF) as implemented in ColabFold was used to generate a predicted ternary complex of a truncated form of Fn and C1r with full-length BBK32. Briefly, the Fn domains 1Fn_1-6_ (Uniprot: P02751, residues 50-345) were co-folded with BBK32 (Uniprot: O50835, residues 21-354). Separately C1r domains CCP1-CCP2-SP (Uniprot: P00736, residues 305-700) was folded with BBK32. The resulting co-folds were structurally aligned to the C-terminal region of BBK32 using Pymol (The PyMOL Molecular Graphics System, Version 3.0 Schrödinger, LLC). Previously resolved crystal structures of the unbound C-terminal region of BBK32 (PDB: 6N1L, [43]), BBK32 in complex with C1r CCP1-CCP2-SP (PDB: 7MZT, [42]), and BBK32 in complex with 1Fn_2-3_ (PDB: 4PZ5, [61]) were used for structural alignments in Pymol to obtain Cα root square mean deviation (RMSD) values.

### Statistics

Data is shown as the average of the replicates with 95% confidence intervals. For antibody-dependent killing assays, two-way ANOVA with a Šidák correction for multiple comparisons was used. All statistical analysis was performed using GraphPad Prism version 9.3.

## RESULTS

### BBK32 Binds Both Fibronectin and C1r Concurrently *In Vitro*

To begin assessing the importance of each of BBK32’s functions during mammalian infection, we sought to determine if the amino-terminal Fn binding and carboxy-terminal C1r interactions were mutually exclusive. Previous work assessing the Fn-binding domain of BBK32 determined that amino acid residues 131-162, inherently disordered until bound by their ligand, were necessary for binding to human fibronectin [47,62]. Residues 206-354 were deemed BBK32’s “ordered” region, forming a stable four helical structure that coordinates the C1r-binding domain [42–44,48]. Because both of BBK32’s activities ostensibly play a role in its pathogenic potential, we were interested in determining how these two functions might affect one another.

We hypothesized that BBK32 might be able to concurrently bind to both Fn and C1r, allowing for both its extracellular matrix binding and complement inhibitory activities during *B. burgdorferi* infection. Using AlphaFold2 (AF2), predictions of each protein-protein interaction were made using full-length BBK32 and relevant domain truncations of each host protein (*i.e.*, 1Fn_1-6_ and C1r CCP1-CCP2-SP) **(Fig. 1)**. The resulting model of the ternary complex was consistent with previously resolved experimental structures in three important ways: i) the C-terminal region of the BBK32 AF2 model closely matches the crystal structure of BBK32-C (RMSD = 0.324 Å; PDB: 6N1L ([43]); ii) the binary complex of the C1r/BBK32 aligns closely with the co-crystal structure of BBK32-C in complex with a proteolytic fragment of C1r (RMSD = 0.831 Å; PDB: 7MTZ ([42]); and iii) similar agreement is found between the binary prediction of Fn/BBK32 and the crystal structure of an N-terminal BBK32 peptide in complex with domains 1Fn_2-3_ of Fn (RMSD = 0.444 Å; PDB: 4PZ5 ([61]).

**Figure 1.**
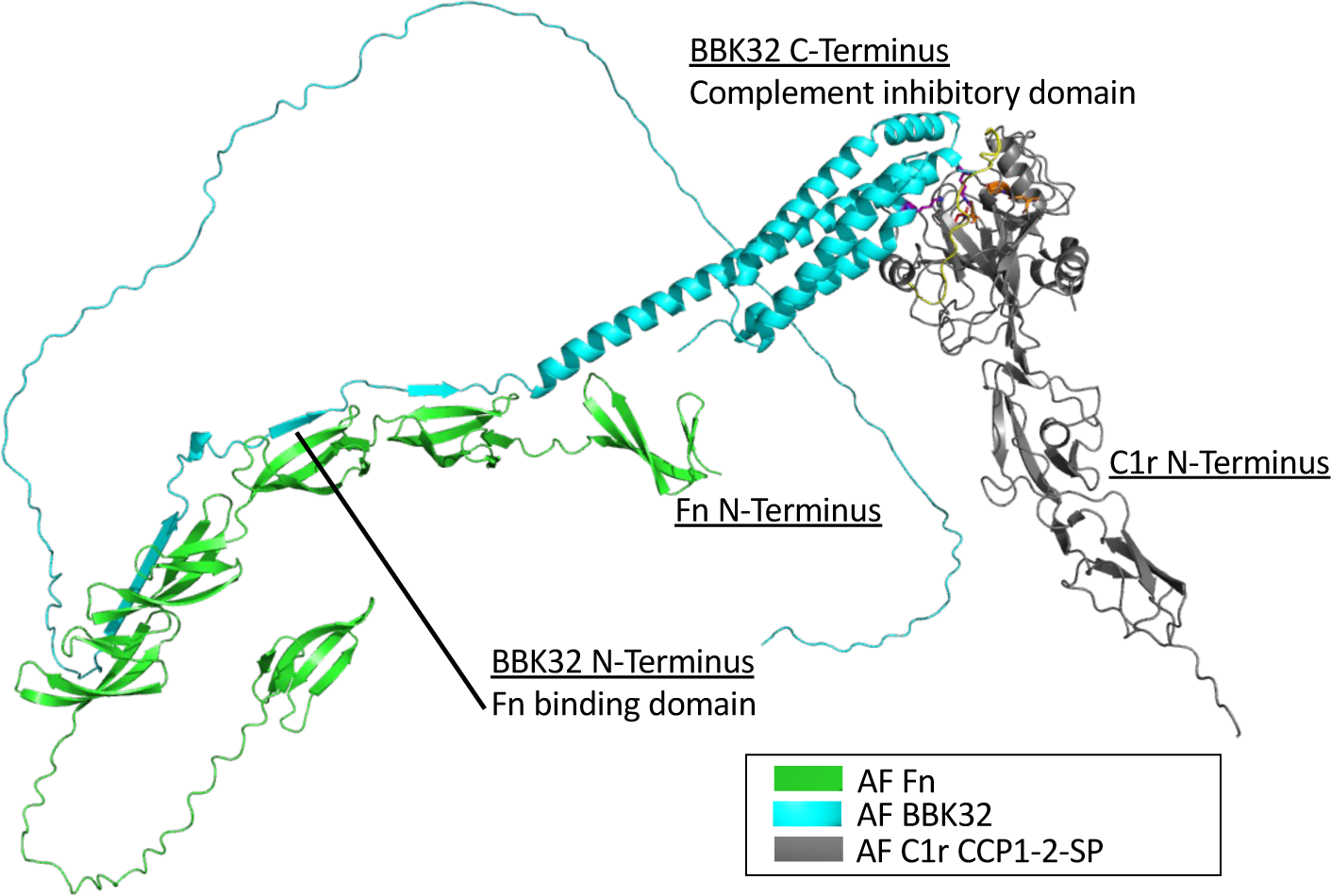
Modeling BBK32’s interaction with C1r and Fn. AlphaFold2 models were used to predict the structure of BBK32 (teal) interfacing with Fn (residues 50-345 [green]) and C1r-CCP1-CCP2-SP (residues 300-705 [gray]).

While the AF2 model presented in Fig. 1 predict that it is structurally feasible for full-length BBK32 to interact with domain fragments of each host ligand. However, given the large size of each full-length host molecule (*i.e.*, Fn (∼500 kDa dimer) and C1 (∼760 kDa as C1qC1r_2_C1s_2_), we sought biophysical evidence for these interactions can occur *in vitro*. Towards this end, we used full-length BBK32 to produce surface plasmon resonance (SPR) biosensors. Purified human C1 (50 nM), purified human Fn (20 nM), or a co-injections of 20 nM Fn and 50 nM C1 were then injected over immobilized BBK32. The resulting sensorgram for 20 nM Fn was then subtracted from the co-injection curve to calculate residual C1 binding by the BBK32 surface **(Fig. 2A-B)**. Analysis of the binding responses strongly suggested that BBK32 can bind both compounds simultaneously *in vitro* **(Fig. 2C)**.

**Figure 2.**
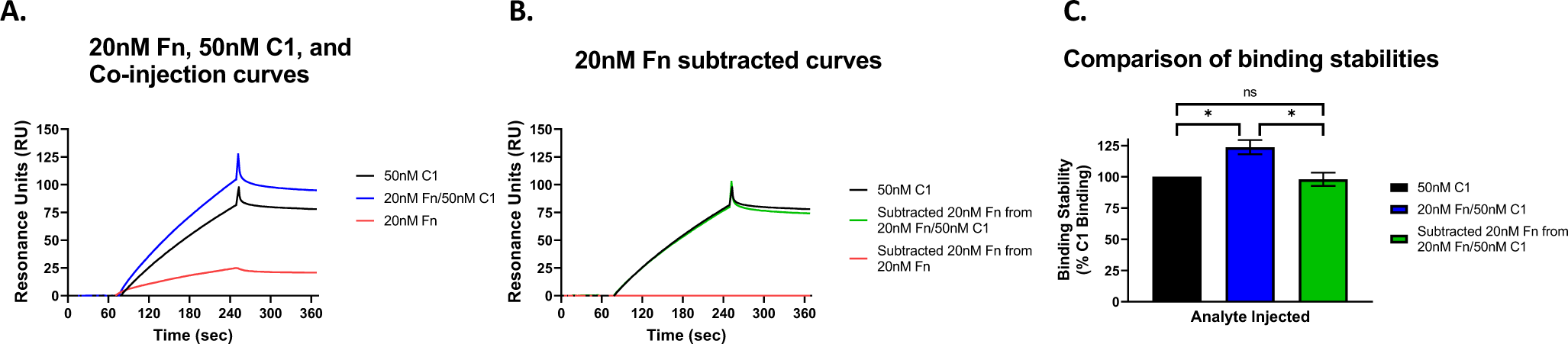
BBK32’s ability to bind both C1 and Fn concurrently. SPR assays were performed using immobilized BBK32 injected with 20 nM human Fn, 50 nM human C1 (A), and a co-injection of 20 nM human Fn and 50 nM human C1 (B). The ability of BBK32 to concurrently interact with human Fn and C1 was calculated by subtracting the human Fn curve from the co-injection curve and comparing that curve to the human C1-alone injection curve (C).

### BBK32 C-terminal double alanine mutant binds Fn but not C1r

We have previously shown that BBK32’s ability to bind and inhibit human C1r is abrogated by double alanine (DA) mutations in its C-terminus. The targets of our prior studies were amino acid residues R248 and K327, which interact with the catalytic and B loop, respectively (also referred to as the K1 and K2 sites recently [41,42]. In order to assess the ability of wild type BBK32 and the DA mutant (BBK32-R248A/K327A) to bind to C1r and Fn, we performed Western blots first showing that protein lysate collected from parent strains (ML23 pBBE22*luc* and B31-A3 GFP), the *bbk32* wild type (WT) complement (JS315 pCD100 and GP100 pCD100), and the *bbk32* DA mutant complement (JS315 pAP7 and GP100 pAP7) produced BBK32, while the *bbk32* knockout strains (JS315 pBBE22*luc* and GP100) did not. We then performed overlays probing these protein lysates with human enzyme C1r and Fn **(Fig. 3A, Fig. S1)**. In these assays, the parent, WT complement, and DA complement bound Fn, while only the parent and WT complement bound C1r, indicating that the DA mutations in the C-terminus did not affect binding of Fn in the N-terminal portion of BBK32. This targeted abrogation of BBK32’s C-terminal C1r binding activity allows for pointed functional analyses of BBK32’s distinct activities. In our work, assays were performed in the ML23 background, previously used in *bbk32* mutant infectivity analyses, for continuity [45,54]. However, unlike the ML23 derivative, the B31-A3 background contains the linear plasmid 25, which is essential to complete the enzootic cycle [54,63]. In addition, screening a distinct *B. burgdorferi bbk32* mutant in an independent infectious derivative provides additional evidence that the results observed were not strain specific. To this end, we used the B31-A3 background **(Fig. 3)** in addition to ML23 **(Fig. S1)** for much of the work, and B31-A3 independently for subsequent competitive overlays, fluorescence microscopy, and flow-based analyses.

**Figure 3.**
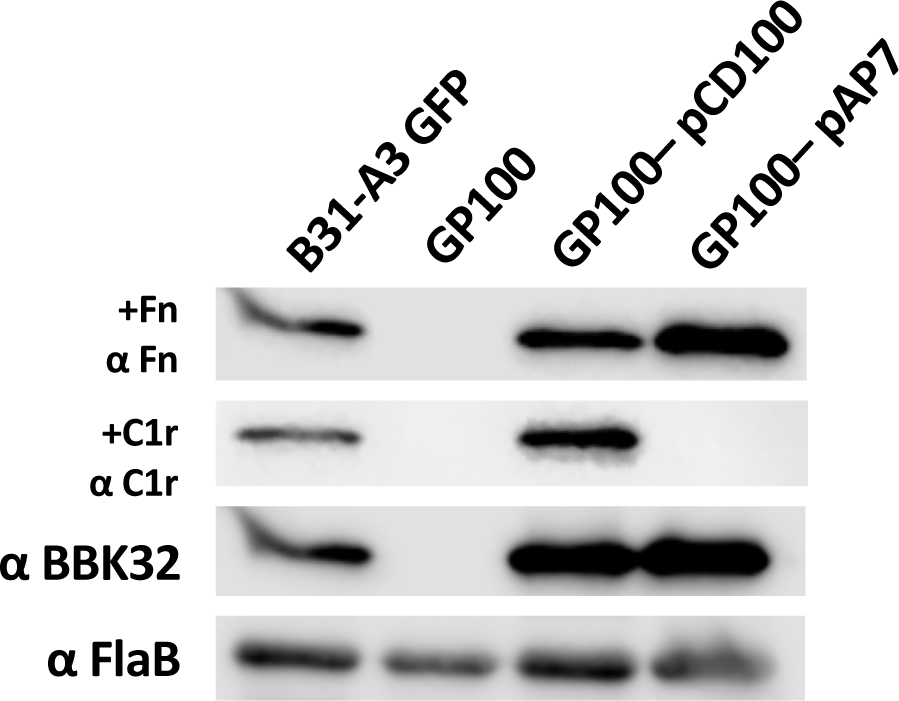
Differential binding of human Fn and C1r to *B. burgdorferi* strain B31-A3, *bbk32* mutant, and functional and non-functional *bbk32* derivatives. Overlay experiments performed using B31-A3 GFP parent, and isogenic *bbk32* mutant strain (GP100), native *bbk32* complement (GP100 pCD100), and a *bbk32-*R248A/K327A double alanine (DA) mutant complement (GP100 pAP7). Protein lysates were probed with human Fn or human C1r and probed with appropriate reagents for detection (see Methods). These same samples were also probed with monoclonal antibodies to BBK32 and *B. burgdorferi* FlaB (third row and bottom row, both panels, respectively).

### BBK32 protects an infectious isolate of *B. burgdorferi* from antibody-mediated complement-dependent killing *in vitro*

Next, we determined the role that BBK32 might play during infection following the development of a *B. burgdorferi*-specific antibody response. We used both the infectious ML23 pBBE22*luc* and B31-A3 background strains for *in vitro* assays in which *B. burgdorferi* were incubated with a targeted polyclonal rabbit antibody against *B. burgdorferi*, coupled, normal human serum (NHS), as a source of complement as performed by Pereira *et al*. [60] Without the addition of the anti-*B. burgdorferi* antibody, both ML23 pBBE22*luc* and B31-A3 are resistant to human serum, as expected based on prior studies determining serum resistance across the *B. burgdorferi sensu stricto* species [17]. As a control for the dependence on borrelial-specific antibody recognition, we used a rabbit isotype antibody control, together with NHS, and observed no killing of the spirochetes **(Fig. 4, Fig. S2)**. However, when all strains were incubated with both NHS and the anti-*B. burgdorferi* antibody, only the derivatives lacking *bbk32* were significantly decreased in their ability to survive antibody-dependent, complement-mediated killing **(Fig. 4, Fig. S2)**. This phenotype was rescued when WT *bbk32* was complemented in the *bbk32* mutant strains, but not in isolates expressing the DA mutated form of BBK32 **(Fig. 4, Fig. S2**). The relative difference between the parent, mutant, and complement strains was comparable in both the ML23 pBBE22*luc* **(Fig. S2)** and B31-A3 **(Fig. 4**) backgrounds, confirming this phenotype is consistent across independently obtained *B. burgdorferi* B31 isolates.

**Figure 4.**
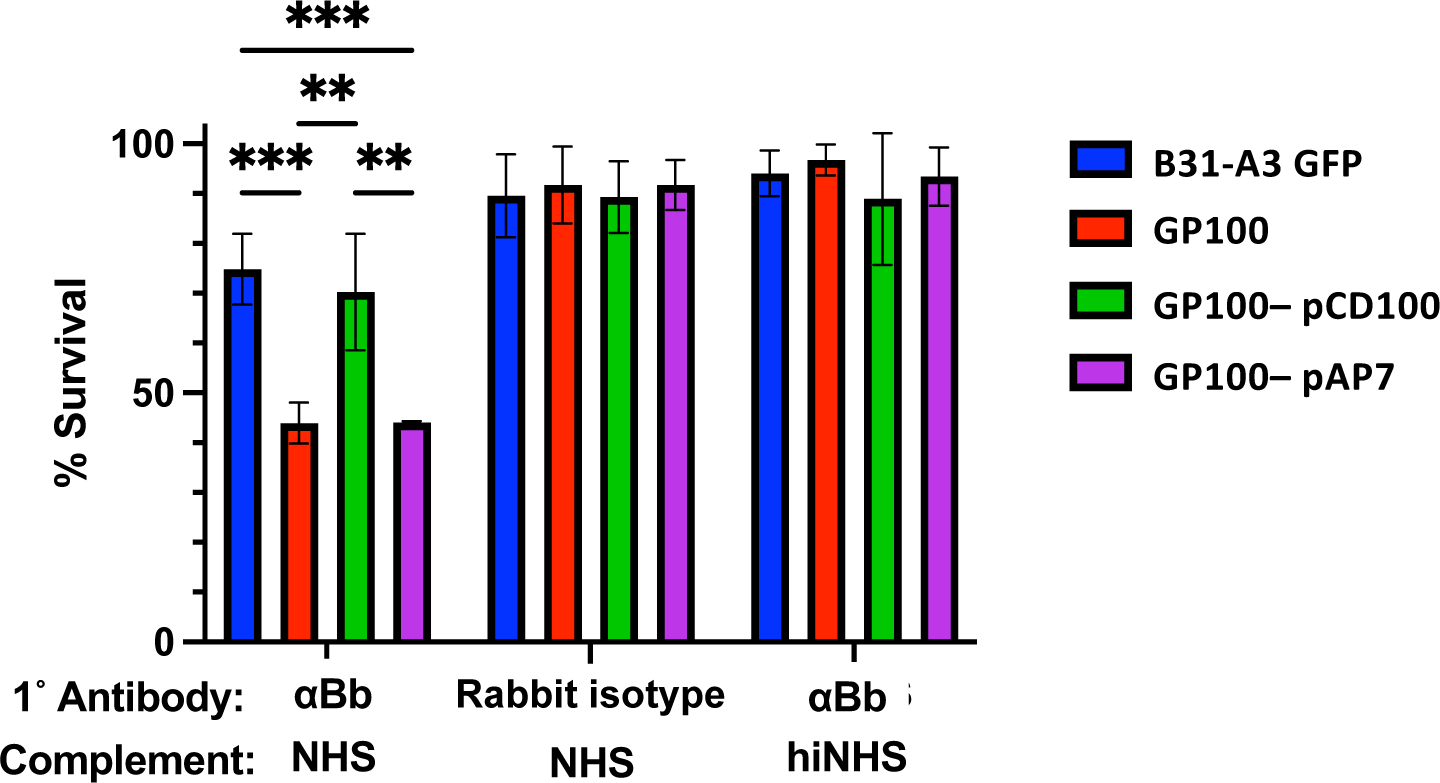
Antibody-dependent complement-mediated killing of *B. burgdorferi* strain B31-A3 derivatives. B31-A3 GFP strains, the isogenic *bbk32* mutant GP100, the GP100 strain complemented with either native *bbk32* (GP100/pCD100) or *bbk32-*R248A/K327A double alanine (DA) mutant (GP100/pAP7) were each separately incubated with anti-*B. burgdorferi* antibody coupled with NHS, rabbit isotype control antibody coupled with NHS, or anti-*B. burgdorferi* antibody coupled with heat inactivated NHS. Viability of each data set was then assessed via dark-field microscopy based on cell motility and overt membrane disruption in triplicate. Error bars represent standard deviation values. ** P <0.01; *** P <0.001.

BBK32 reduces complement activation on the surface of *B. burgdorferi*.

To further depict sensitivity to antibody-dependent complement-mediated killing in *bbk32* deficient strains, immunofluorescence was used to qualitatively assess whether functional BBK32 could inhibit the classical complement cascade. As our readout, surface deposition of C4c—as a proxy for C4—was scored following incubation with antibodies specific for *B. burgdorferi* and human serum. To the end, C5-depleted serum was used to prevent degradation of *B. burgdorferi* cells via formation of the MAC during processing of cells. The targeted cells remain intact in the C5-depleted serum as the absence of C5 prevents the generation of the MAC but, importantly, retain the upstream modification of complement markers involved in complement activation, here C4c. As such, these cells are subject to complement activation steps upstream of C5 cleavage without any further downstream effects. The parent strains and *bbk32* mutants that expressed WT *bbk32* had reduced C4c localized on their surface. In contrast, *B. burgdorferi* cells lacking BBK32, and particularly for those expressing the *bbk32* DA mutant in a *bbk32* mutant background, showed more C4c—as a proxy for complement activation—when incubated with the anti-*B. burgdorferi* antibody (**Fig. 5**). When incubated with a rabbit isotype control antibody and the C5-depleted serum, all cells, independent of genetic composition, showed a decrease in both antibody binding and less C4c deposition consistent with reduced complement activation (data not shown).

**Figure 5.**
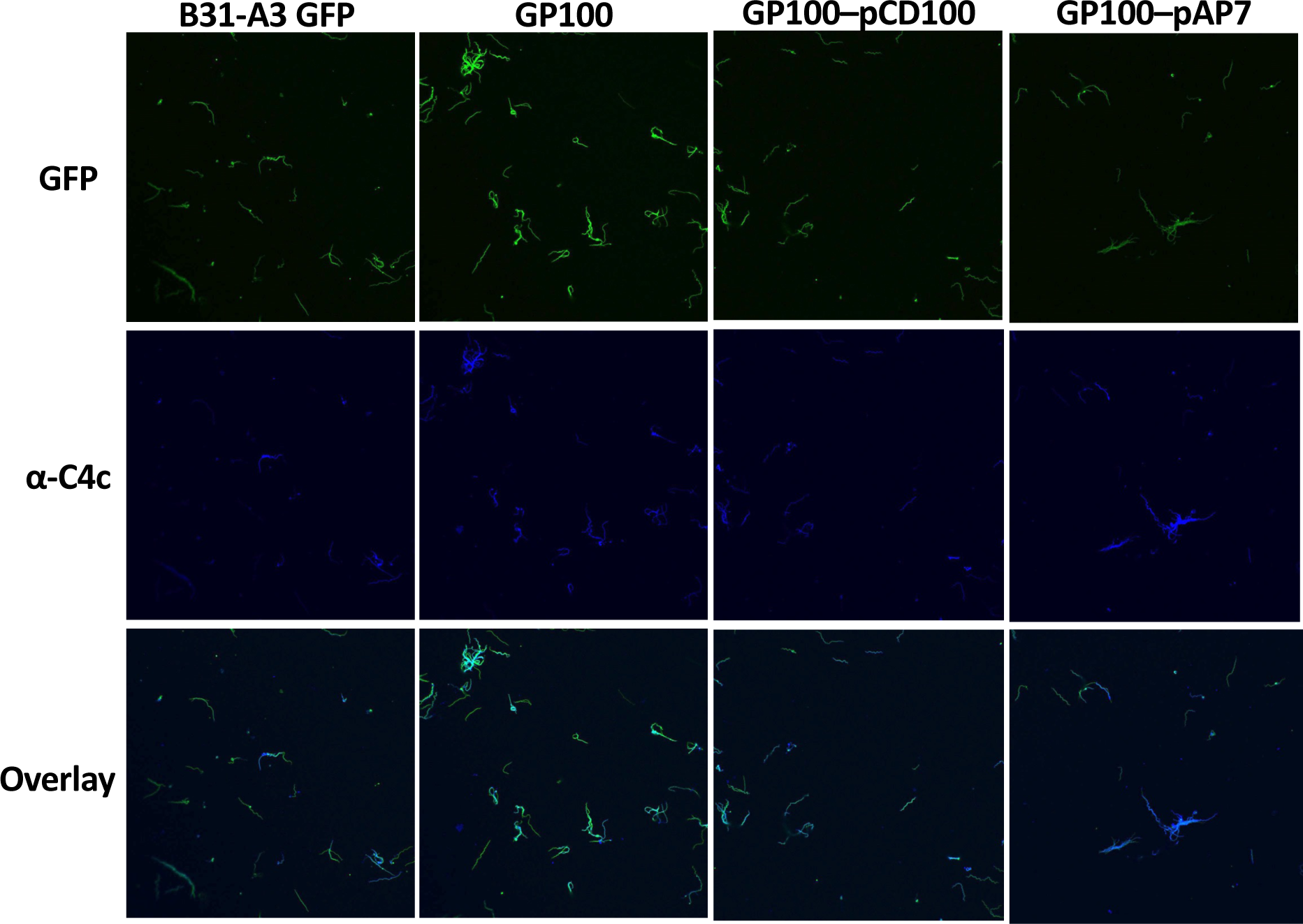
Qualitative assessment of increased deposition of complement components in *bbk32* mutant strains. B31-A3 GFP (A), GP100 (B31-A3 GFP Δ*bbk32* (B), GP100 pCD100 (Δ*bbk32* with native *bbk32* complement) (C), and GP100 pAP7 (Δ*bbk32* with native *bbk32* DA complement) (D) were incubated with rabbit anti-*B. burgdorferi* antibody coupled with C5-depleted serum. Cells were then probed with murine anti-C4c, followed by anti-rabbit and anti-mouse Alexa Fluor 594 and Cy5, respectively. Cells were fluorescently imaged via confocal microscopy, and colocalization of the anti-*B. burgdorferi* antibody and C4c was assessed.

To address the deposition of C4c in a quantitative manner, we employed flow cytometry of the of the *B. burgdorferi* parent and *bbk32* mutant strains. The results demonstrated that the parental borrelial strain had limited C4 on its surface (**Fig. 6A**). In contrast, the *bbk32* mutant exhibited more C4 deposited consistent with functional BBK32 inhibiting C1r and reducing C4 proteolysis and subsequent attachment of C4c to the borrelial cell surface (**Fig. 6B**). While not statistically significant, the trend indicates a reduction in C4c deposition in cells that have functional BBK32 (**Fig. 6C**). Taken together, these results show an antibody dependence for complement-mediated killing of infectious *B. burgdorferi in vitro* and demonstrate a role for BBK32 in protection from complement-based clearance. This BBK32 activity is likely to be important during established infection when a potent *B. burgdorferi* humoral response is present.

**Figure 6.**
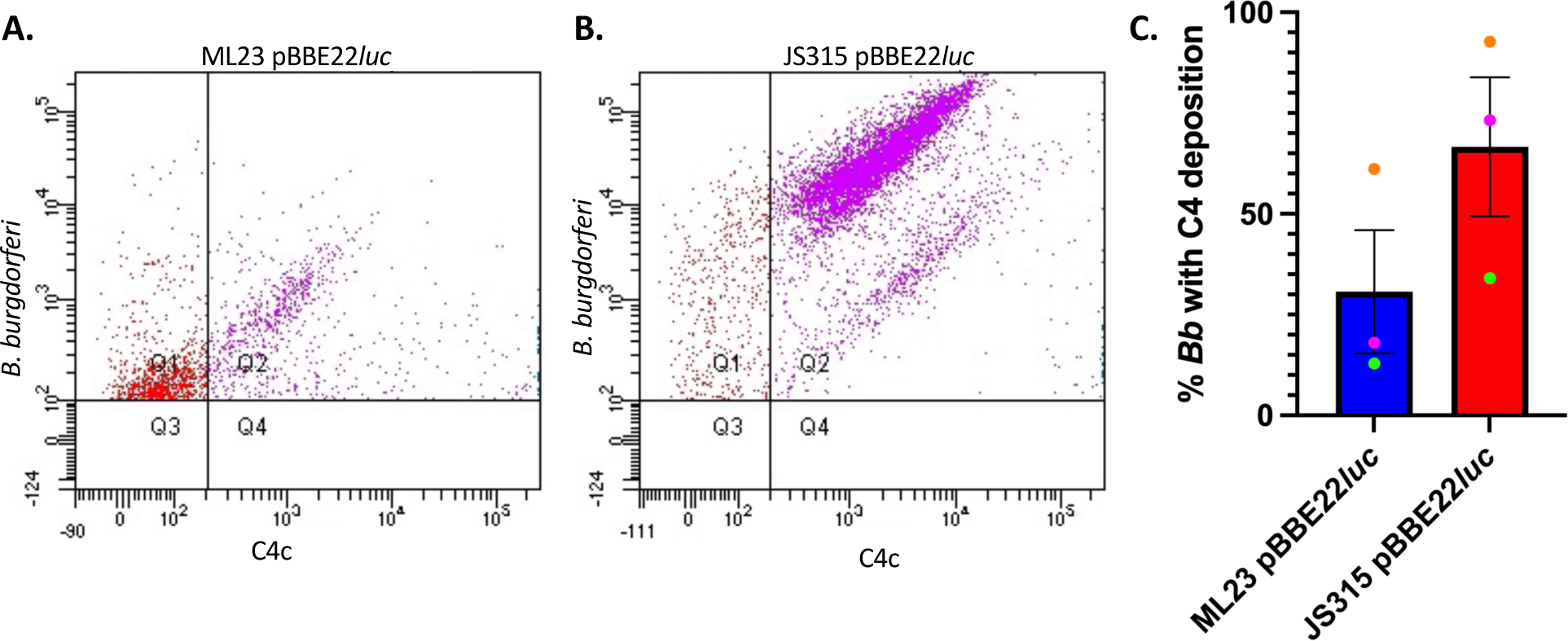
Flow cytometry scores for increased C4 deposition in borrelial cells lacking *bbk32*. Parental *B. burgdorferi*, along with an isogenic *bbk32*, were incubated with polyclonal antibodies against *B. burgdorferi* with C5-depleted human serum. The level of C4c deposition was then scored for 10,000 events using flow cytometry. The scatter plots are shown for the parent strain (**A**) and the *bbk32* mutant (**B**) from a single representative analysis. The extent of C4 on the surface was scored based on the florescence associated with the Cy5 secondary antibody against a C4c monoclonal antibody. The percent of the parental *B. burgdorferi* strain positive for C4c deposition relative to the *bbk32* mutant was compared for three individual replicates, indicated by the green, pink, and orange data points, for 10,000 events each (**C**). Shown are the standard errors for three individual replicates.

## DISCUSSION

BBK32 is known to contribute to *B. burgdorferi*’s ability to establish experimental infection, thought in part to be due to its ability to bind ECM components and, more recently, its ability to inhibit complement activation [43,44,46–50,53]. The temporal balance of these two functions is unknown, and whether *B. burgdorferi* is forced to prioritize interacting with one of BBK32’s ligands during the dissemination phase of infection, where extravasation and complement inhibition are both crucial, has not been explored. Though the impact of normal human serum on a non-infectious isolate of *B. burgdorferi* had been characterized in our previous work [42], the survival of an infectious derivative of *B. burgdorferi* in the presence of a borrelial-specific antibody had not been assessed. We hypothesized that survival of infectious *B. burgdorferi* following exposure to antibody against borrelial antigens-- and subsequent activation of classical complement--would be diminished by the presence of functional BBK32.

As an extracellular pathogen, *B. burgdorferi* is found in blood and interstitial fluid that contain innate immune compounds and cells. Early in infection, and as the infection proceeds, some *B. burgdorferi* are processed by the host, resulting in a potent antibody response against them. Despite a strong humoral response to *B. burgdorferi,* as well as the presence of host complement, *B. burgdorferi* persists, in part, by quelling classical pathway activation despite borrelial antibody-antigen complex formation [24,64]. One additional question that has not been explored is whether BBK32 can bind the C1 complex and ECM components (namely Fn) simultaneously. Our analysis here of infectious isolates with fully functional BBK32, or a BBK32 mutant deficient in classical complement inhibition, is designed to serve as an *in vitro* proxy for testing this possibility. The work described herein portrays BBK32’s C1r-inhibitory activity in a novel context: protection against targeted antibody-dependent complement killing, which occurs concurrent with BBK32’s Fn-binding activity. The use of the impaired *bbk32* DA complement afforded us a unique strategy for characterization of a multifunctional protein in *B. burgdorferi.* Namely, normal Fn binding but with an abrogated C1r interaction. Herein, we correlated native *bbk32* expression with decreased localization of complement components to the *B. burgdorferi* surface and showed the protective impact of BBK32 against antibody-directed complement-mediated killing. This analysis provides the foundation for pointed structure-guided mechanistic characterization of other C1r inhibitors across Lyme disease and tick-borne relapsing fever (TBRF) spirochetes [41,57].

Orthologues to *bbk32* have been identified in TBRF *Borrelia*, some of which have already been determined to bind both Fn and complement components similar to BBK32 [41,57,65]. By leveraging our understanding of the amino- and carboxy-terminal activities of these proteins, we can begin to ask how these multifunctional domains contribute to infection in the face of natural and targeted antibody responses. *B. miyamotoi* FbpA is an example of a C1r inhibitor in a TBRF spirochete that also interfaces with Fn [57]. Through the identification of key “hotspot” residues that mediate BBK32’s and *B. miyamotoi* FbpA’s interactions with C1r, we have been able to define a consistent, species-agnostic mechanism for this class of complement inhibitors for a subset of TBRF FbpA proteins [41,57] Interestingly, independent of their ability to bind Fn, all the C1r inhibitor proteins characterized to date—either BBK32, BBK32-like, or Fbp proteins—contain a disordered N-terminal domain and an ordered C-terminal domain, the latter binding to and mediating the inhibition of C1r [42–44,57] Mutations that abrogate C1r binding and inhibition allow for a pointed analysis of Fn binding in *B. burgdorferi* pathogenesis could be leveraged for determining the role of the distinct domains found within BBK32. In this study, we have elucidated the ability of BBK32 to bind to both C1r and Fn concurrently. However, a targeted mutagenesis approach is needed to assess how these domains affect borrelial pathogenesis. While we already have mutations in BBK32 that abrogate C1r binding and inhibition, we lack an analogous construct that reduces or eliminates BBK32-mediated Fn binding. To this end, future studies will aim to mutagenize BBK32’s ECM-binding domain in key residues for its interaction with soluble or insoluble Fn [47,48,61,62] to determine if Fn-and/or C1r-binding activities promote pathogenic outcomes associated with borrelial infection.

There are several pathogens, notably *Staphylococcus aureus*, that produce surface proteins that interact with host ECM and complement components [62,66–68]. This common dual function feature of ECM and complement interaction by additional pathogenic bacteria suggests that this correlation between pathogens is not random and is involved in mediating their pathogenic potential [69,70] The idea that both distinct functional domains of these proteins might be contributing to immune evasion seems plausible, especially considering Fn’s immunomodulatory activity through its ability to bind C1q’s collagen-like tail. This Fn::C1q interaction is thought, in some instances, to alter the phagocytosis of pathogens and promote their recognition by phagocytes [71–73]. As this relates to *B. burgdorferi*, BBK32 does not interact with Fn in regions thought to be recognized by phagocytes; thus, it is unlikely that it functions to prevent Fn-mediated phagocytosis by occluding this site. However, BBK32 could be positioning Fn to interact with C1q’s collagen tails and sequestering C1q in complex with Fn, limiting the recognition of the C1 complex by C1q receptors. This type of beneficial immune evasion tactic might also explain BBK32’s ability to bind to the C1r zymogen, which previously seemed counterintuitive to the goal of preventing C1r/C1 complex localization to the *B. burgdorferi* surface [42,44,71,72]. Nonetheless, this theory would need to be further investigated.

It is known that infectious *B. burgdorferi* is resistant to human complement due to the presence of numerous genes that encode proteins that inhibit the various complement pathways [26–28,74]. The best characterized of these proteins are the factor H binding proteins that inhibit C3 convertase formation and provide resistance to the alternative pathway of complement [31,32,75–78] Given that BBK32 is an inhibitor of the classical complement cascade, we tested *bbk32* mutants relative to the infectious parent strain for their ability to bind to immobilized C1. We found that there were no differences between these two strains, implying that there were compensatory borrelial proteins that could also bind C1 and potentially inhibit the classical complement pathway [44]. Recently, Pereira et al. used a lipoprotein library to identify novel proteins that are capable of binding to C1 [60] In addition to BBK32, the cp32-encoded Elp proteins recognized C1 [60]. The borrelial genome encodes at least five Elp paralogues that share 44-59% identity and 59-76% similarity. These proteins, namely ElpB, ElpD, ElpM, ElpQ, and ElpX, are encoded by *elp* genes with flanking sequences that are nearly identical. As such, the use of conventional allelic exchange to inactivate all *elp* genes will be difficult. This is needed ostensibly to determine how classical complement evasion factors into borrelial infectivity relative to BBK32. To address this limitation, several CRISPR-based modalities could be used to either knock down or inactivate *elp* genes with or without intact *bbk32* [79–82] We are currently pursuing these approaches.

The work described in this study was limited to *in vitro* analyses that tracked the role of BBK32 in survival of *B. burgdorferi* via the classical pathway. All the assays used employed borrelial antibodies and active classical complement component to assess survival using in vitro killing assays (**Fig. 4**), as well as C4 deposition by fluorescence microscopy (**Fig. 5**) and flow cytometry (**Fig. 6**). The data obtained indicates that functional BBK32 is needed for enhanced survival under these conditions. For most of these approaches the complementation of the *bbk32* mutant with shuttle vectors was achieved with intact *bbk32* sequence, but not when the carboxy terminal mutations in *bbk32* (e.g., *bbk32*R248A/K327A) were present. Following up on these phenotypes *in vivo* is crucial to deciphering the role that each of the BBK32 forms contribute to infection. Attempts to address infectivity were made, and while the known attenuated phenotype of the *bbk32* mutant was retained, complementation was not achieved, presumably due to inadequate *in vivo* selection of the *trans* complemented *bbk32* alleles. In addition, another confounding variable is the presence of other C1-inhibitors, such as the recently described Elp proteins that also inhibit the classical pathway [60,83,84]. Thus, isolating mutants in multiple genes simultaneously may be necessary to observe a reduced infectivity phenotype, at least in the context of classical complement inhibition. With the growing number of complement-inhibitory genes identified in borrelial species, this approach is timely, and future studies evaluating the functions of complement inhibitors with multiple interactions should consider a potential for interplay between the distinct activities. As noted above, the advent of CRISPR-Cas9 systems facilitates approaches to simultaneously knock down or inactivate multiple targets in various combinations [79–82] These studies should define which borrelial complement inhibitory protein(s) are essential for borrelial pathogenic readouts.

In summary, we utilized a *bbk32* mutant and a strain expressing a *bbk32* allele deficient in C1r inhibition to assess the spatial dynamics of BBK32’s interaction with ECM and complement components. We also demonstrated that BBK32 confers protection from antibody-initiated complement-mediated killing using whole cell sensitivity assays, microscopy, and flow cytometry as readouts. Regarding antibody-dependent, complement-mediated killing, BBK32 could serve to quell the deleterious effect of antigen-antibody interactions that increase as the infection proceeds and class switching occurs. Through our previous structure-guided analyses that identified key residues of BBK32 that interface with C1r [42], we used biochemical analyses to determine BBK32’s ability to bind Fn and the C1 complex concurrently *in vitro*, and cell-based experiments to assess these residues’ role in antibody-dependent complement protection. This work also describes a strategy that could be applied to other multifunctional proteins in the borrelial C1r-inhibitor group. Taken together, these studies contribute to our understanding of borrelial surface-expressed proteins capable of interacting with distinct host targets and provide a framework for characterization of TBRF C1r inhibitors orthologous to BBK32.

## Notes

### Competing Interest Statement

The authors have declared no competing interest.

