## Supplemental Data for "The dual activity of BBK32: Implications for simultaneous inhibition of borrelial-specific antibody-dependent complement activation and fibronectin binding"

**SUPPLEMENTAL FIGURE LEGENDS**

**Figure S1. Differential binding of human Fn and C1r in *B. burgdorferi* and *bbk32* mutant ML23 derivatives.** Overlay experiments were performed using the ML23 pBBE22*luc* parent, a *bbk32* mutant strain (JS315 pBBE22*luc*, native *bbk32* complement strain (JS315 pCD100), and a *bbk32*-R248A/K327A double alanine (DA) mutant complement (JS315 pAP7). Protein lysates were incubated with either human Fn or human C1r and probed with appropriate reagents for detection (see Methods). These same samples were also probed with monoclonal antibodies to BBK32 and *B. burgdorferi* FlaB (third row and bottom row, both panels, respectively).

**Figure S2. Antibody-dependent complement-mediated killing of *B. burgdorferi* of strain B31 ML23 derivatives.** ML23 pBBE22*luc*, the isogenic *bbk32* mutant JS315, the JS315 strain complemented with native *bbk32* (JS315/pCD100) and a *bbk32*-R248A/K327A double alanine (DA) mutant complement (JS315 pAP7) were each separately incubated with anti-*B. burgdorferi* antibody coupled with NHS, rabbit isotype control antibody coupled with NHS, or anti-*B. burgdorferi* antibody coupled with heat inactivated NHS. Viability was then assessed via dark-field microscopy based on cell motility and overt membrane disruption in triplicate. Error bars represent standard deviation values. * P <0.05; *** P <0.001.

**SUPPLEMENTAL FIGURES**

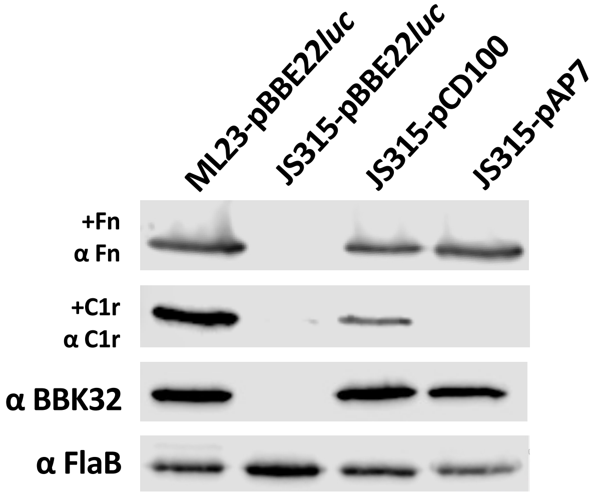


**Figure S1.**

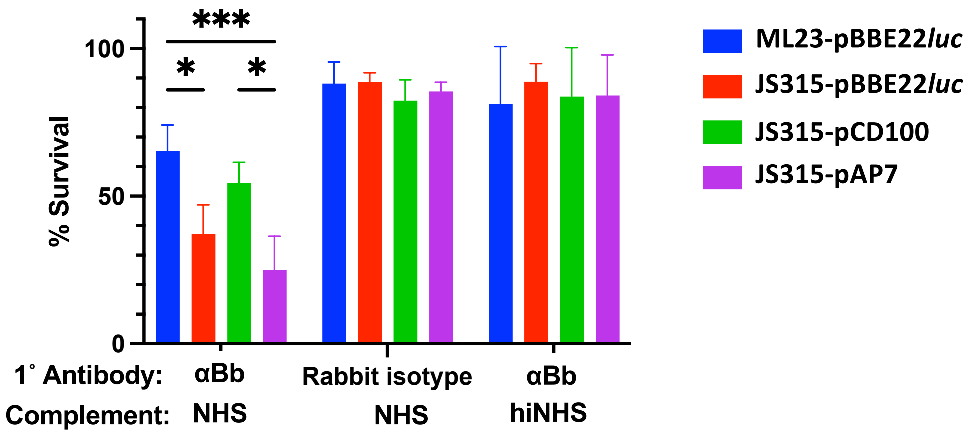
***
*
*
100
50
0
% Survival
1˚ Antibody:
Complement:
αBb
NHS
Rabbit isotype
NHS
αBb
hiNHS
ML23-pBBE22*luc*
JS315-pBBE22*luc*
JS315-pCD100
JS315-pAP7

**Figure S2**
